# The absence of F-box motif Encoding Gene SAF1 and Chromatin Associated factor CTF8 together contributes to MMS Resistant and HU Sensitive phenotype in S. *cerevisiae*

**DOI:** 10.1101/656363

**Authors:** Meenu Sharma, V. Verma, Narendra K Bairwa

## Abstract

The Replication factor-C compex which related to cohesion, constitutes, three subunits called Ctf18, Ctf8 and Dcc1. These three subunit complex assist the loading of PCNA onto the chromosome. None of the replication factor C components are essential for cell viability. The null mutant of the CTF8 in *S.cerevisiae* shows the chromosome instability and high frequency of chromosome loss. The SAF1 gene product of S. *cerevisiae* involved in the degradation of adenine deaminase factor Aah1p by SCF-E3 ligase, which itself is the part of E3 ligase. The ubiquitin marked degradation of Aah1p occurs during nutrient stress which lead to cell enter into the quiescent state. The N-terminus of Saf1p interacts with the Skp1 of SCF-E3 ligase and at C-terminus recruits with Aah1p. Here we have investigated about the binary genetic interaction between the SAF1 and CTF8 genes. The strains containing single and double gene deletions of SAF1 and CTF8 were constructed in the BY4741 genetic background. Further the mutant strains were evaluated for growth fitness, genome stability and response to genotoxic stress caused by hydroxyurea (HU) and methyl methane sulfonate (MMS). The *saf1Δctf8Δ* strain showed the increased growth phenotype in comparison to *saf1Δ, ctf8Δ, and* WT strain on YPD medium. However *saf1Δctf8Δ* strain when grown in the presence MMS showed resistance and HU sensitive phenotype when compared with *saf1Δ, ctf8Δ.* The frequency of Ty1 retro-transposition was also elevated in *saf1Δctf8Δ* in comparison to either *saf1Δ* or *ctf8Δ.* The number of cells showing the two or multi-nuclei phenotype was also increased in *saf1Δctf8Δ* cells when compared with the either *saf1Δ* or *ctf8Δ.* Based on these observations, we report that the absence of both the gene SAF1 and CTF8 together leads to MMS resistance, HU sensitivity, and genome instability. This report warrants the investigation of mechanisms of differential growth phenotype due to loss of SAF1 and CTF8 together in presence of genotoxic stress in future.

## Introduction

The ubiquitin proteasome system regulates the phase transition due to nutrient availability in *Saccharomyces cerevisiae* (FINLEY *et al.* 1987).The nutrient deprivation condition induces stress, which leads to cell enter into the quiescent phase (FINLEY *et al.* 1987).The Saf1, of *S.cerevisiae* which constitutes the SCF E3-ligase component, recruits the Aah1p, for proteasomal mediated degradation during nutrient deprivation conditions (ESCUSA *et al.* 2006; ESCUSA *et al.* 2007). The Aah1, adenine deaminase of *S.cerevisiae*, converts adenine to hypoxanthine, implicating the role of Saf1 in nucleotide metabolism. The genetic interaction studies reported negative interaction of SAF1 with CDC10, CDC11, CDC12, and HYP2 (COSTANZO *et al.* 2016). The null mutant of SAF1 showed the synthetic growth defects with HSP82 (ZHAO *et al.* 2005), POL2 (DUBARRY *et al.* 2015) and RTT109 (FILLINGHAM *et al.* 2008) RRM3 (Sharma, *et al.* 2019, bioxrv archived data).

SAF1 also showed the negative genetic interaction with RFC-Ctf18 complex member DCC1 (COSTANZO *et al.* 2016). RFC helps in loading of PCNA (a processivity factor), known as sliding clamp, onto the DNA. The PCNA clamps the DNA polymerase onto its template (PETRONCZKI *et al.* 2004).The replicative RF-C helps in switching between DNA polymerases (delta and epsilon) during lagging strand DNA synthesis (BYLUND and BURGERS 2005).There are four types of RF-Cs (SHIOMI *et al.* 2007).The RF-C which related to cohesion, constitutes, three subunits called Ctf8, Ctf18, and Dcc1 proteins (BERMUDEZ *et al.* 2003). None of these proteins are essential for viability. The absence of RF-C complex showed the cohesion defects. Ctf8 forms the part of a complex (Ctf18p, Ctf8p, Dcc1p) called RFC which help in sister chromatid cohesion and chromosome transmission (MAYER *et al.* 2004).The deletion of any member of the complex individually or in combination causes cell to be sensitive to microtubule depolymerizing drugs and showed sister chromatid cohesion defects (MAYER *et al.* 2004). The complex help in loading of proliferating cell nuclear antigen (PCNA) onto primed or gapped DNA in ATP dependent manner. The loading of PCNA is mediated by the single-stranded DNA binding protein RPA (BYLUND and BURGERS 2005). The complex fails to load PCNA onto nicked or single-stranded circular DNA. There is evolutionary functional homologue complex reported in the nucleus of human cells carrying out the similar function (MERKLE *et al.* 2003). One of the Ctf18-RFC complex components, Ctf8 was first discovered as a mutant who showed the decreased chromosome transmission fidelity in mitosis in a haploid yeast strains with normal sensitivity to ultraviolet and gamma-irradiation (SPENCER *et al.* 1990). The null mutant of CTF8 in yeast showed the severe chromosome instability and high frequency of chromosome loss. The mutant showed lower competitive fitness, slow growth, abnormal bud morphology (WATANABE *et al.* 2009) and increased sensitivity to multiple chemicals, including MMS, hydroxyurea. Synthetic genetic arrays (SGA) study with the CTF8 deletion reported the role of other genes such as CHL1, CSM3, BIM1, KAR3, TOF1, CTF4, and VIK1 in sister chromatid cohesion (MAYER *et al.* 2004). The RFC, Ctf18 has been proposed to be candidate cancer drug target in a synthetic lethal genetic interaction study carried out in *Caenorhabditis elegans* (MCLELLAN et al. 2009). The complex also involved in preserving genome stability through the suppression of the triplet repeat instability which is a hallmark of the hereditary neurological disorders (GELLON *et* al. 2011; OKIMOTO *et al.* 2016).The mutants of the Ctf18-RFC complex showed the triplet repeat instability which included expansions, contractions, and fragility.

Ty1 are the retrotransposons element which constitutes the part of yeast genome. They are flanked by long terminal repeats and contains genomic feature which encodes structural components of the virus like particles and the reverse transcriptase (BOEKE *et al.* 1985; GARFINKEL *et al.* 1985). The Ty1 element remains dormant in wild type cells due to host encoded factors and transpose infrequently due to inactivity of the repressive factors (SCHOLES *et* al. 2001). The transposition events are considered to be mutagenic hence indicator of the defect in the genome maintenance.

The duplicated nuclei need to be segregated accurately in the all eukaryotic cells for precise distribution of genetic material. In *Saccharomyces cerevisiae*, nuclear migration from mother cell to daughter cell after duplication involves moving of nucleus closer to budneck then insertion of elongating nucleus into the daughter cell during anaphase (COTTINGHAM and HOYT 1997). The migration of nucleus from mother to daughter cells depends on the action of cytoplasmic microtubules (CARMINATI and STEARNS 1997). Mutations in tubulin affects the cytoplasmic microtubules which affects the nuclear migration and causes abnormal nuclear division within the mother cells (HUFFAKER *et al.* 1988). The abnormal nuclear division leads to accumulation of two or more nuclei within the mother cells which could lead to genome instability.

The genetic interaction between SAF1 gene and RFC-complex member CTF8 has not been investigated so far. Here, we report the binary genetic interaction between the two genes SAF1 and CTF8. The simultaneous deletion of both genes (SAF1 and CTF8) leads to increased growth phenotype and elevated frequency of Ty1 retro-transposition. The nuclei in the double mutant showed the nuclear migration defects. The double mutant also exhibited the resistance to MMS and sensitivity to HU in comparison to single gene mutant or WT.

## EXPERIMENTAL PROCEDURES

### Yeast Strains and plasmid

The yeast strains with their genotypes and plasmids used in this study are mentioned in the **Table 1** and **Table 2.**

**Table 1:**
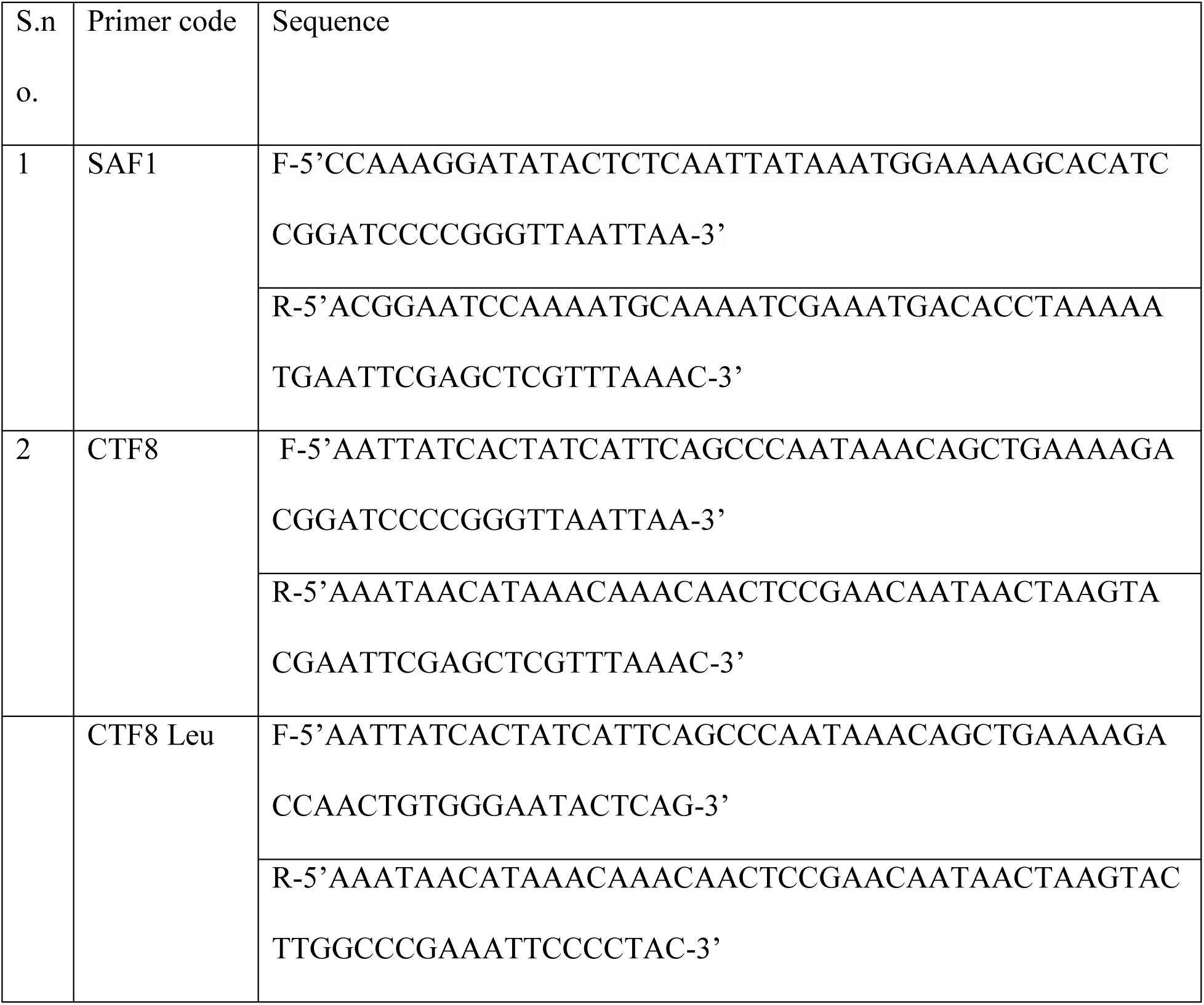
List of primers used for construction of deletion strains.

**Table 2:**
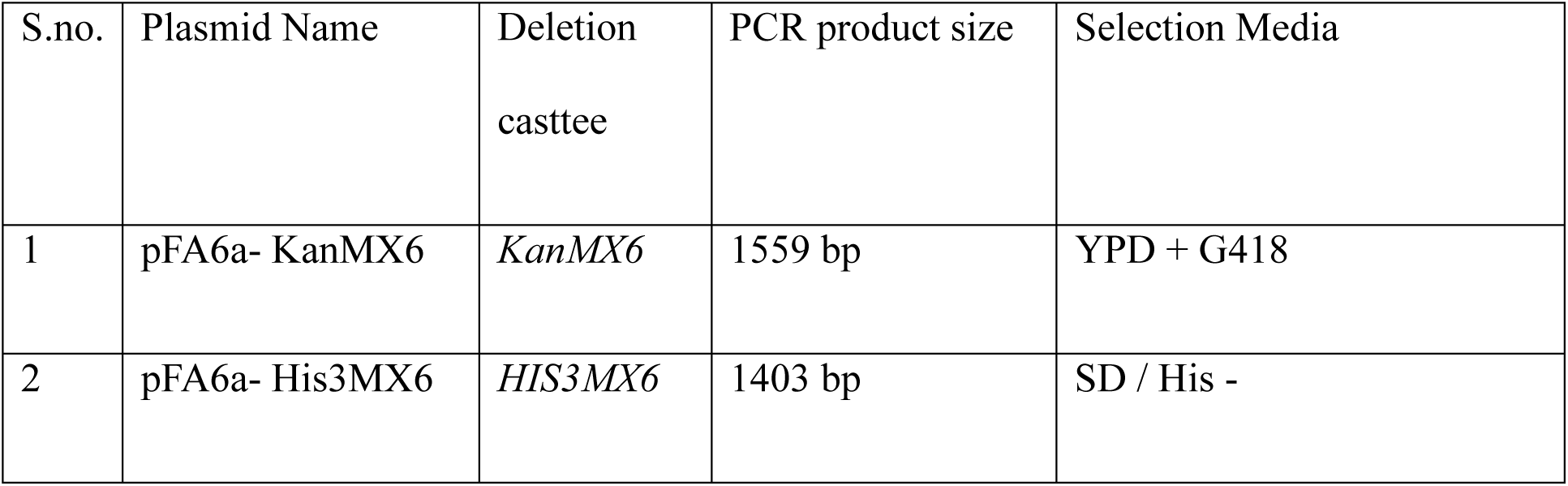
List of plasmids used for generating deletion cassette.

| S.no. | Plasmid Name | Deletion<br>casttee | PCR product size | Selection Media |
| --- | --- | --- | --- | --- |
| 1 | pFA6a- KanMX6 | *KanMX6* | 1559 bp | YPD + G418 |
| 2 | pFA6a- His3MX6 | *HIS3MX6* | 1403 bp | SD / His - |

**Table 3:** Yeast strains and their genotype used in the study.

| S.no. | Strain | Genotype | source |
| --- | --- | --- | --- |
| 1 | BY4741 | MATa his3Δ1 leu2Δ0 met15Δ0 ura3Δ0 | Dr. Deepak Sharma<br>IMTECH |
| 2 | *saf1Δ* | *saf1Δ::HIS3* | This study |
| 3 | *ctf8Δ* | *ctf8Δ::KanMX* | This study |
| 4 | *saf1Δctf8Δ* | *saf1Δ::HIS3, ctf8Δ::KanMX* | This study |
| 5 | JC2326 | *MAT-ura3, cir0, ura3-167, leu::hisG, his32*<br>*Ty1his3AI-270, Ty1NEO-588, Ty1 (tyb::lacZ)-146* | Prof. M. Joan Curcio<br>USA |
| 6 | *saf1Δ* | *saf1Δ::KanMX* | This study |
| 7 | *ctf8Δ* | *ctf8Δ::KanMX* | This study |
| 8 | *saf1Δctf8Δ* | *saf1Δ::KanMX, ctf8::LEU2* | This study |

### Growth Assay

For growth assessment of the WT and mutants in YPD broth and on solid meadia the method mentioned in (Sharma *et al.* 2019, biorxiv achived data; doi 1101/636902, www.biorxiv.org) was adopted. Briefly, The growth of strains in broth was measured at 2 hrs of interval for 14 hrs. The OD of three independent replicates of each culture at every time point was measured and average of that was plotted against the time. Growth of the strains was also compared by streaking on the YPD plates followed by incubation at 30°C for 2-3 days.

### Phase Contrast Microscopy

To compare the morphology of WT and deletion strain, phase contrast microscopy was conducted using Leica DM3000 microscope at 100X magnification. Each strain was grown up to log phase in YPD medium at 30°C and imaged.

### Spot Assay

To compare the growth fitness of WT and mutants in presence of genotoxic agents such as hydroxy urea (HU) and methyl methane sulfonate (MMS) the method was followed as mentioned in (Sharma *et al.* 2019, biorxiv achived data; doi 1101/636902, www.biorxiv.org). Briefly, wild type (BY4741) and its deletion derivatives strains (*saf1Δ, ctf8Δ,* and *saf1Δctf8Δ*) were grown in the 25 ml YPD (Yeast Extract 1% w/v, Peptone 2% w/v, dextrose 2% w/v) medium overnight at 30°C. The next day the cultures were diluted and grown in fresh YPD medium for 3-4 hrs so as to reach log phase (OD600 0.8-1.0). The cultures were equalized by OD at 600nm. A ten-fold serially diluted sample (3µl) was spotted onto agar plates containing (YPD and YPD + HU and MMS). The plates were incubated at 30°C for 2-3 days and imaged.

### Assay for Ty1 retro-mobility

To measure the retro-mobility of Ty1 element methos mentioned in (SCHOLES *et al.* 2001; BAIRWA *et al.* 2011) and (Sharma *et al.* 2019, biorxiv achived data; doi 1101/636902, www.biorxiv.org) was followed. Briefly, the assay was carried out using JC2326 as WT and deletion derivatives (*saf1Δ, ctf8Δ,* and *saf1Δctf8Δ).* Each strain carries a single Ty1 element which is marked by an indicator gene HIS3AI. The reporter gene HIS3 is split by artificial intron (AI) in an orientation opposite to the HIS3. When Ty1 which is marked with this arrangement undergoes transcription, splicing followed reverse transcription to generated cDNA copy and finally integration into genomic locations. The integration of the cDNA would generate HIS3 positive phototrophs. The quantitative measurement of the HIS3 positive colonies is considered as frequency of retro-mobility. To measure the Ty1 retro-transposition frequency in the WT (JC2326; reporter strain) and the deletion derivatives (*saf1Δ* and *ctf8Δ* and *saf1Δctf8Δ*), a single colony of each strains was inoculated into 10 ml YPD broth and grown overnight at 30°C. The overnight grown cultures were again inoculated in 5 ml YPD at 1:1000 dilutions. The cultures were allowed to grow up to saturation point (144hrs) at 20°C. The saturated culture was serially diluted and plated on minimal media (SD/His^-^ plates) followed by incubation at 30°C for 3-7 days. The frequency of appearance of HIS^+^ colonies was measured for Ty1 retro-mobility.

### Assay for Nuclear Migration defects using Fluorescence microscopy

For detection of nuclear migration defects, assay described by (PALMER *et al.* 1992; BRACHAT *et al.* 1998) and (Sharma *et al.* 2019, biorxiv achived data; doi 1101/636902, www.biorxiv.org) was adopted. Briefly, strains were grown to early log phase (OD_600_ ∼ 0.8) at 30°C. Yeast cells were washed with distilled water and suspended in 1X PBS (Phosphate Buffer Saline). Further, fixation was done by addition of 70% ethanol before DAPI staining. Cells were washed with 1X PBS then again centrifuged for 1 minute at 2500 rpm. DAPI stain (1mg/ml stock) to final concentration of 2.5µg/ml was added and incubated for 5 minutes at room temperature and visualized under UV light of fluorescent microscope with 100X magnification. A total of 200 cells were counted and grouped according 0, 1, 2, and multi nuclei per cell, more than two nuclei per cell indicated the nuclear migration defect.

### Statistical methods

Statistical significance of observations was determined using paired student t-test. P-value less than 0.05 indicated significant.

## RESULTS

### Deletion of both the genes SAF1 and CTF8 together leads to growth advantage

The deletion of an ORF may compromise the growth fitness of the mutant depending on whether the gene is essential or non-essential. The essential gene cannot be deleted however the deletion of non-essential gene does not affect the growth fitness on the rich medium to a large extent. Both the gene SAF1 and CTF8, the non-essential therefore the null mutant of the SAF1 and CTF8 was reported to be viable (GIAEVER *et al.* 2002). We wished to determine the impact of deletion of both the gene together on growth fitness in rich medium. We constructed the single (*saf1*Δ, *ctf8*Δ) and double gene deletion (*saf1*Δ*ctf8*Δ) in the BY4741 genetic background and assessed the growth in the YPD broth and solid medium. We observed that single gene mutant (*saf1*Δ, *ctf8*Δ) were viable. The growth of the single gene mutants (*saf1*Δ, *ctf8*Δ*) was* slightly reduced in comparison to WT. In contrast, the double mutant (*saf1*Δ*ctf8*Δ) growth was observed better than the WT (**Figure 1 A, C**). The phase contrast imaging of the single gene mutant (*saf1*Δ, *ctf8*Δ) appeared normal. However, the (*saf1*Δ, *ctf8*Δ) mutant showed the slight enlargement of the cell size (**Figure1B**). The *saf1*Δ*ctf8*Δ showed the synthetic growth advantage as indicated by streaking and growth kinetics.

**Figure 1.**
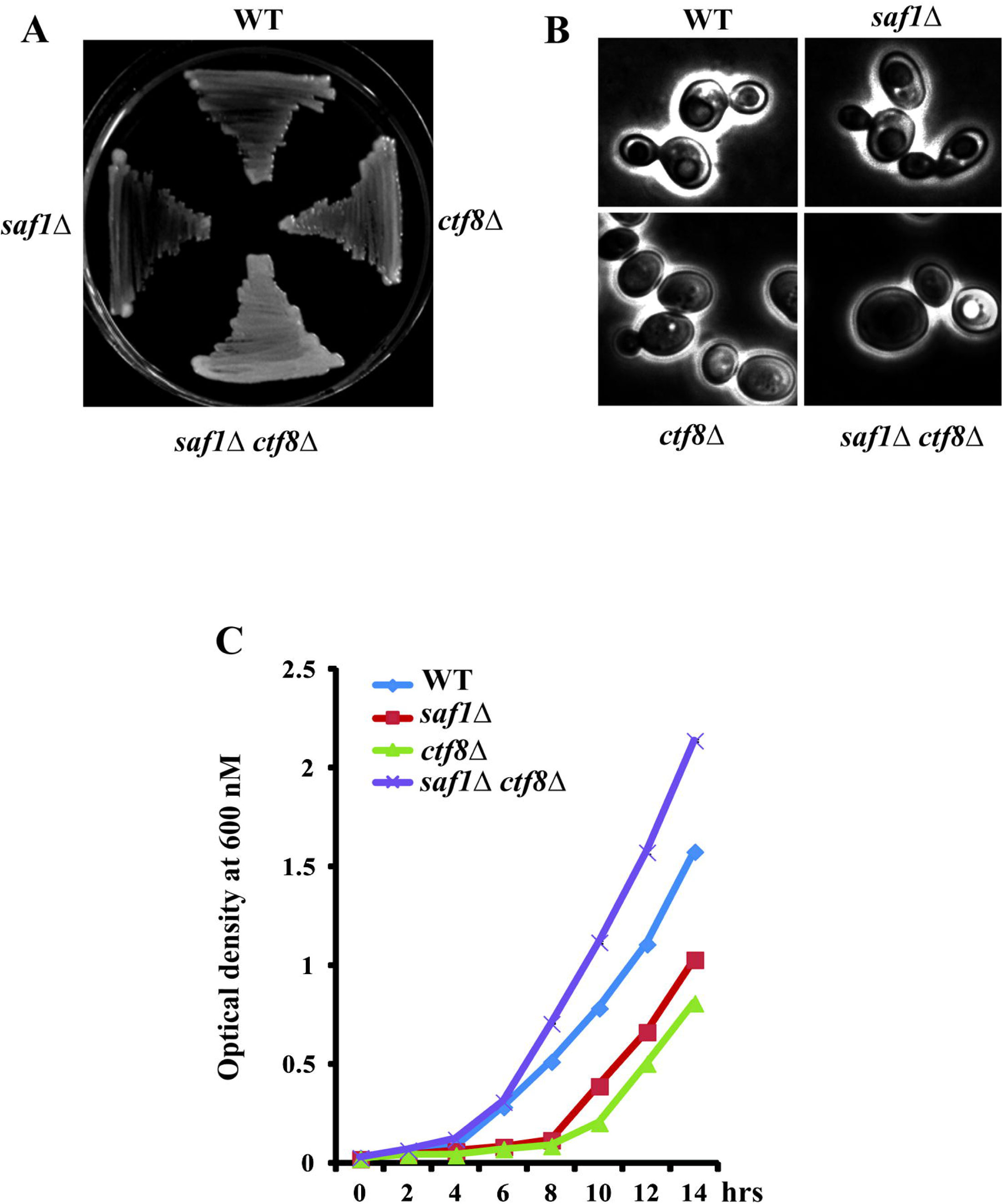
Comparative analysis of growth and morphology of WT, *saf1*Δ, *ctf8*Δ, *saf1*Δ*ctf8*Δ strains. A growth of streaked strain on YPD plates which were incubated for 2 days at 30°C and then photographed. **B.** Phase contrast images of log phase cultures at 100X magnification using Leica DM3000. **C**. Growth Kinetics of strains (WT, *saf1*Δ, *ctf8*Δ, *saf1*Δ*ctf8*Δ). Cells were collected every 2 hour period and cellular growth was measured by optical density (OD) at 600 nm using TOSHVIN UV-1800 SHIMADZU. The data shown represent the average of three independent experiments. The error bars seen represent the standard deviation for each set of data.

### Absence of both the gene SAF1 and CTF8 together leads to MMS resistance and HU sensitivity

The genotoxic stress causing agents such as hydroxyurea and MMS have been utilized for mutant screen and clinical use. The hydroxyurea (HU) act as replication checkpoint inhibitor reduces the dNTP pool in the cell. The mechanism involves the inhibition of the ribonucleotide reductase (RNR) activity. We wished to determine the growth response of WT and mutants (*saf1*Δ, *ctf8*Δ, and *saf1*Δ*ctf8*Δ) in the presence of 200mM HU by semi-quantitative spot analysis. Previously it has been reported that *ctf8*Δ cells showed sensitive phenotype in presence of hydroxyurea. In spot assay, WT, *saf1*Δ, *ctf8*Δ appeared slightly sensitive *to* HU however *saf1*Δ*ctf8*Δ cells showed the increased sensitivity to HU when compared with the growth on YPD agar medium without the HU (**Figure 2A**). However the *saf1*Δ*ctf8*Δ *cells showed the* resistance phenotype in comparison to WT or single gene mutant when grown on YPD agar plates containing 0.035% MMS **(Figure 2B**). Further, mechanism of DNA damage repair need to be investigated in *saf1*Δ*ctf8*Δ *cells* treated with MMS.

**Figure 2.**
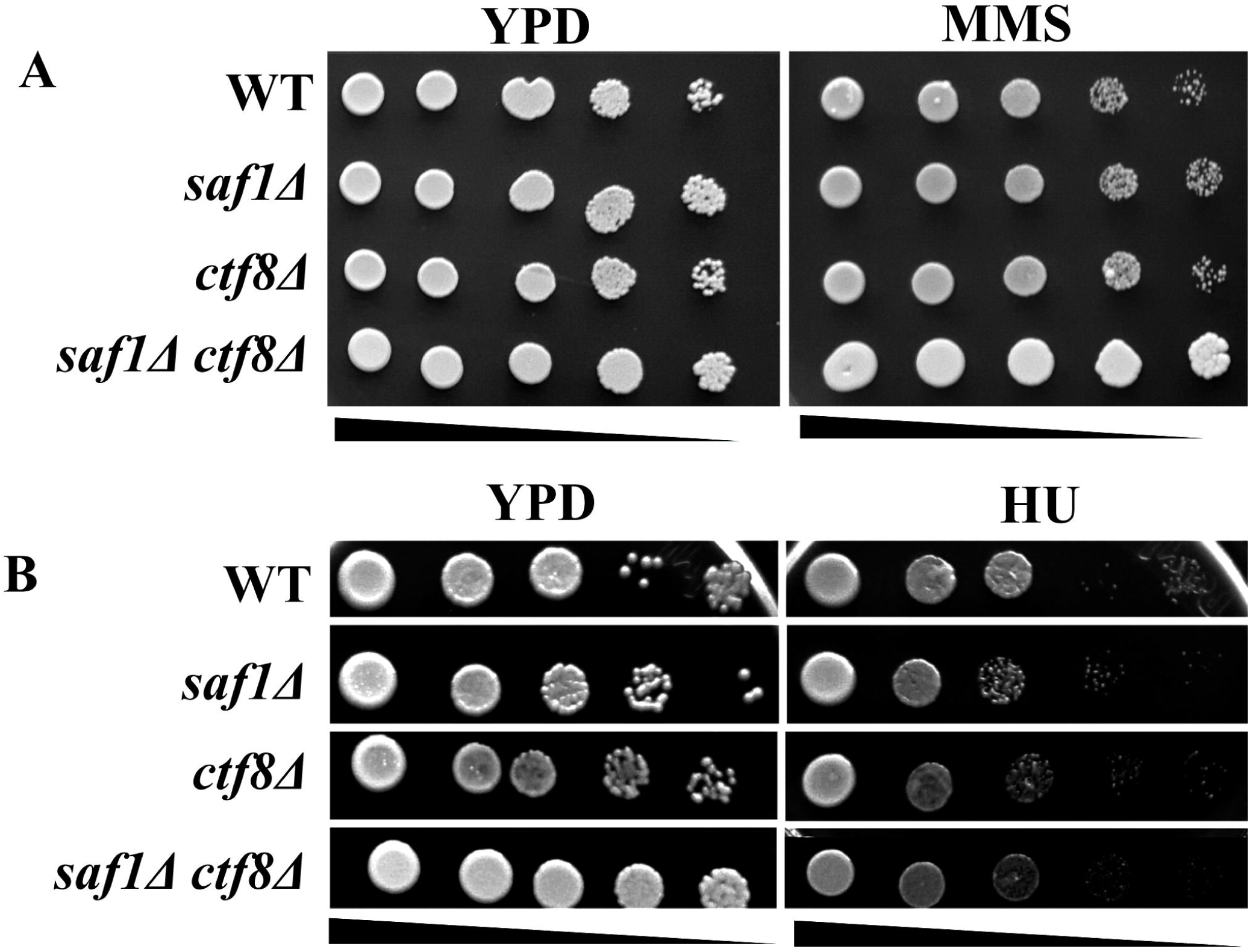
Comparative assessment of growth phenotype of WT *saf1*Δ, *ctf8*Δ, *saf1*Δ*ctf8*Δ strains in presence of methyl methane sulfonate and hydroxyurea by spot analysis. Each strains was grown to log phase was equalized by O.D 600nm and ten fold serially dilution was made. From each dilution 3µl spotted on YPD, YPD + HU/ MMS containing agar plates. The *saf1*Δ*ctf4*Δ showed the resistacne to MMS and sensitivity to HU

### SAF1 and CTF8 together contributes to nuclear migration defects

The transfer of the newly replicated nuclear DNA into the daughter cells is very delicate process. The DAPI stain which is very specific to the nuclear DNA allows the visualization of the DNA in the form single nuclei in case of WT cells having intact DNA or the multi-nuclei which may indicate the defects in the nuclear migration from mother to daughter cells. We wished to investigate the status of nuclear DNA migration in saf1Δ, *ctf8*Δ, *saf1*Δ*ctf8*Δ by staining of the log phase culture with DAPI staining. The stained cells were visualized under the fluorescence microscope using appropriate filter. We observed that WT cells showed the compact nuclei (**Figure 3 A)** and majority of cells (80%) showed single nuclei representing budded cells, in case of *saf1*Δ percentage of cell showing the multinuclei phenotype was increased (**Figure 3 B**). However *ctf8*Δ showed increased percentage (20%) of cells showing multi-nuclei phenotype. However *saf1*Δ*ctf8*Δ cells showed nuclear DNA at the bud neck junction (**Figure 3A**) and multi nuclei phenotype in nearly 29% of the cells (**Figure 3B**). The multi-nuclei phenotype indicated the defect in the nuclear migration. The data indicates both the gene SAF1 and CTF8 together contributes to accuarate nuclear distribution between mother and daughter cell however mechanism need to investigated.

**Figure 3.**
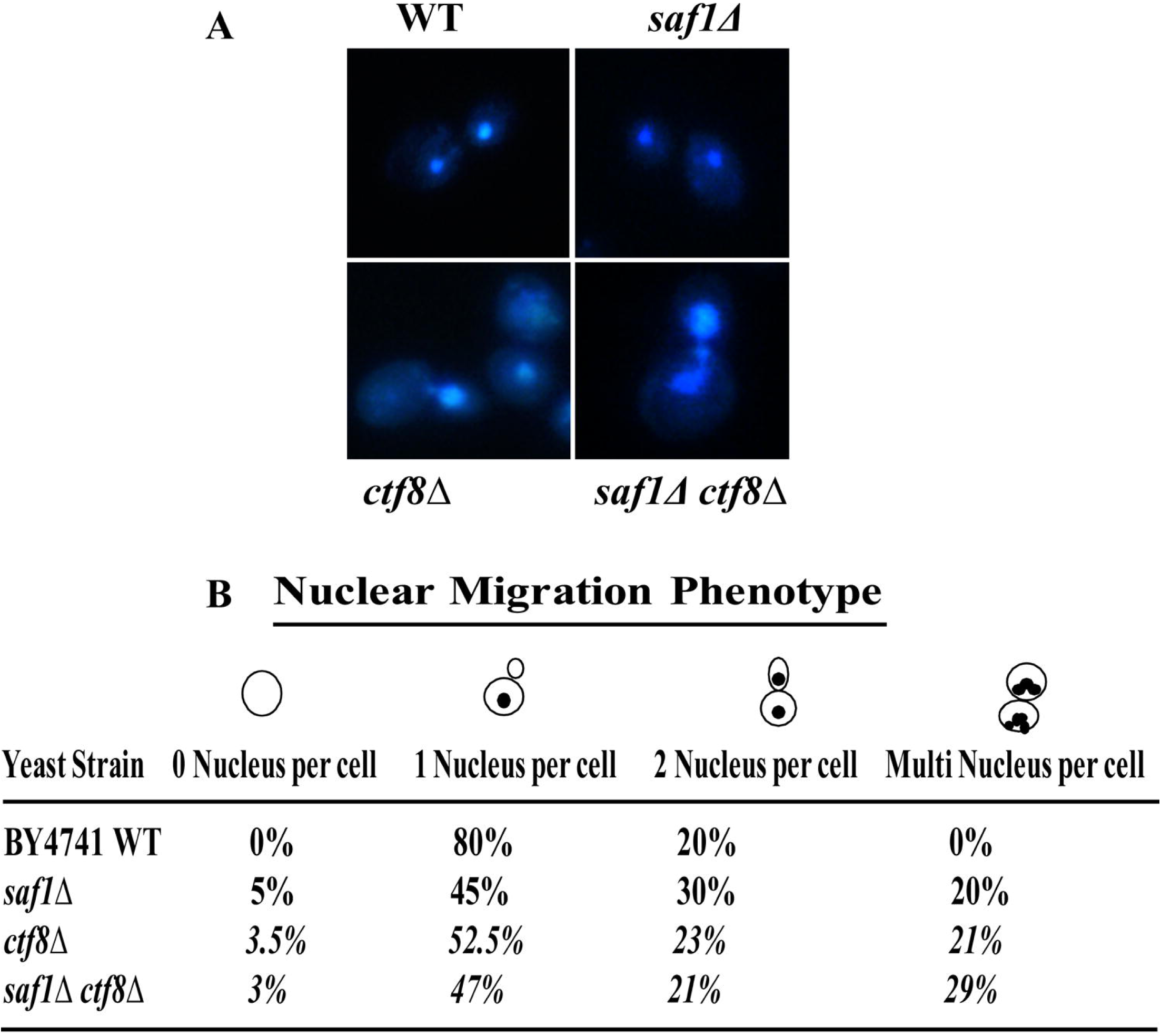
Assessment of nuclear migration phenotype using DAPI (4′,6-diamidino-2-phenylindole) staining of WT, *saf1*Δ, *rrm3*Δ, *saf1*Δ*ctf8*Δ. **A** Representative fluorescent images of WT, *saf1*Δ, *ctf8*Δ, *saf1*Δ*ctf8*Δ cells showing the status of Nuclear DNA migration. The images were acquired at the 100X magnification using Leica DM3000 fluorescent microscope. B Table showing the percentage from the count of 200 cell as, 0, 1, 2 and multi-nucleus in each strains, more than two nuclei indicate the nuclear migration defect..

### Deletion of both SAF1 and CTF8 induced high frequency of Ty1 retro–transposition

Replication stress and DNA damage known to induce the Ty retro-mobility in the S. *cerevisiae.* Ctf8 constitutes the part of RFC-Ctf18 complex which helps in the loading of PCNA. The PCNA clamps the DNA polymerases epsilon onto replicating legging strand. We wished to find out whether the absence of both genes could impact the genome stability as their absence showed the faster growth phenotype in comparison to WT. The increase in the frequency of Ty1 retro-transposition is an indicator of the genome instability phenotype. Previously the highest frequency of Ty retro-transposition reported in the *rrm3*Δ*tof1*Δ background was 550 fold above the WT level(BAIRWA *et al.* 2011). We measured theTy1 retro-mobility in the WT, saf1Δ, *ctf8*Δ *and saf1*Δ*ctf8*Δ cells using the previously described split HIS3 reporter assay, which allowed the counting of HIS+ phototrophs formation on the SD/his-plates. Using this assay we observed that WT and *ctf8*Δcells did not show the detectable level of His+ phototrophs formation however the *saf1*Δ cell showed the nearly 8-10 fold increase in the Ty1 retro-transposition in comparison to WT (**Figure 4 A, B**), however when both the mutant combined *(saf1*Δ*ctf8*Δ*) showed* nearly 170 fold increase in the Ty1 retro-transposition **(Figure 4A, B**). However the mechanism of increased Ty1 retro transposition needs further investigation in future.

**Figure 4.**
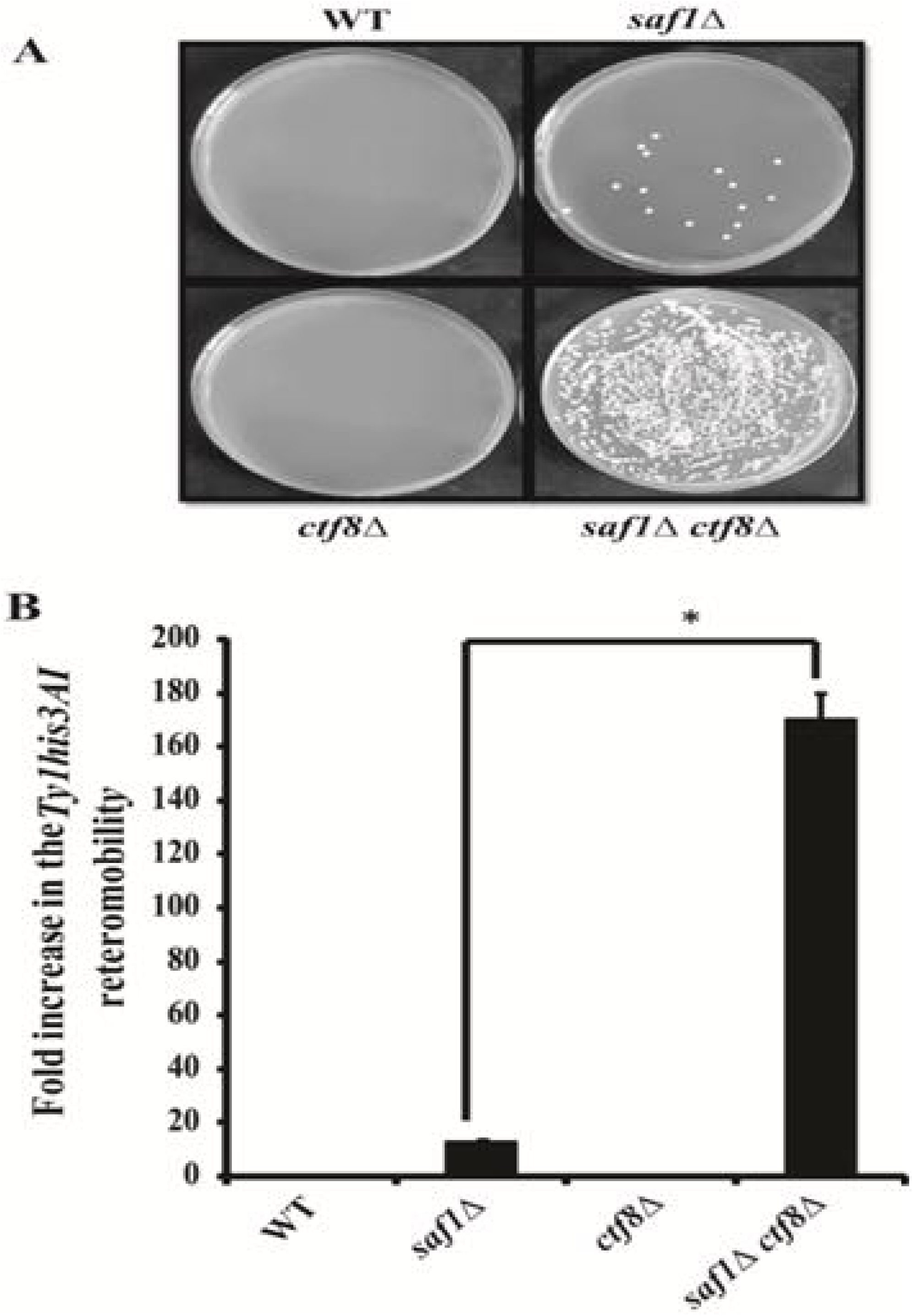
Assay for genome instability by measuring of HIS*3AI* marked Ty1 transposition frequency in WT, *saf1*Δ, *ctf8*Δ, *saf1*Δ*ctf8*Δ. A Images of plates showing the Ty1 transposition induced colonies on SD plate lacking His media. B Bar diagram showing the frequency of Ty1his3AI transposition in each strains. The data shown represent the average of three independent experiments. The significance of transposition was determined by using two tailed t-test. P - value (p) less than 0.05 indicate significant difference and the symbol * represent to p<0.05.

## Discussion

In this paper we have investigated the genetic interaction between the SAF1 and chromosome associated genome stability regulator CTF8. The search for the genetic interactions and their impact on phenotype and biological processes is important for the precision medicine development. The investigations on the binary interactions are crucial for better understanding of the system biology and gene networks. This work reports that SAF1 and CTF8 genes together regulate the growth fitness and loss of both leads to the additive growth in rich medium and resistance to genotoxic agent MMS and sensitivity to HU. The Dcc1 component of the Rfc18 complex (Ctf18, CTf8, and Dcc1) reported to have negative genetic interaction in a high-throughput study (COSTANZO *et al.* 2016). Previously the *ctf8*Δ cell showed the decreased growth in the presence of MMS (CHANG *et al.* 2002) and HU (HARTMAN and TIPPERY 2004; PARSONS *et* al. 2004). The *saf1*Δcells showed the resistance to histone deacetylase inhibitor CG-1521 drug(GAUPEL *et al.* 2014). However the loss of both the genes SAF1 and CTF8 together showed the resistance to MMS and sensitivity to HU suggesting their functioning in different pathways. This is the first report on the genetic interactions between the SAF1 and CTF8 gene which showed positive interactions during normal growth conditions. It would interest to investigate the trigenic interaction among the SAF1, CTF8 and DCC1 and impact on the growth fitness in the presence of HU and MMS. How the loss of genes affects the biological process is the next step in understanding of the overall system biology. Further the data presented in this study on Ty1 retro transposition suggest that loss of either CTF8 or SAF1, elicit the weak retro transposition event. However deletion of both the gene together leads to increased Ty1 retro-transpositions, indicating genome instability phenotype. The absence of both the gene together also showed the increased percentage of cells showing the nuclear migration defects, thereby suggesting the role of SAF1 and CTF8 interaction in faithful segregation of the duplicated genome from mother to daughter cells. We propose that the increased growth phenotype in the absence of both the gene leads to increase in the genome instability and drug resistance which coincide with the cancer cell phenotype. The double mutant (*saf1*Δ*ctf8*Δ) can be used for screening of the chemical compound. It also be interesting to investigate the RNAi mediated knockdown of the hCTF8 and HERC2 in the human cell for fast growth phenotype and resistance to genotoxic agents.

## Acknowledgment

We thank to Prof. M. Joan Curio, Dr. Deepak Sharma, IMTECH, Dr. Ravi Manjithya, NCBS, strains and plasmids. We also thank Dr. Jitendra Thakur, NIPGR, Dr. Preeti Sharma, Mr. Pervez S. Slathia, SMVDU for use of their equipment facility.

## Funding information

This work was supported by a grant (BT/RLF/Re-entry/40/2012) from the Department of Biotechnology, GOI, and New Delhi to N.K.B who is recipient of the Ramalingaswami fellowship from DBT, New Delhi.

## Conflict of Interest

The authors declare that they have no conflicts of interest with the content of this article.

## Author’s contributions

NKB conceived and directed the study and wrote the paper with MS and VV. MS performed the experiments and analysed with NKB. VV provided the bioinformatics facility and analysed the data. All the authors reviewed the results and approved the final version of manuscript.

## References

Bairwa, N. K., B. K. Mohanty, R. Stamenova, M. J. Curcio and D. Bastia, 2011 The intra-S phase checkpoint protein Tof1 collaborates with the helicase Rrm3 and the F-box protein Dia2 to maintain genome stability in Saccharomyces cerevisiae. J Biol Chem 286: 2445–2454.

Bermudez, V. P., Y. Maniwa, I. Tappin, K. Ozato, K. Yokomori et al., 2003 The alternative Ctf18-Dcc1-Ctf8-replication factor C complex required for sister chromatid cohesion loads proliferating cell nuclear antigen onto DNA. Proc Natl Acad Sci U S A 100: 10237–10242.

Boeke, J. D., D. J. Garfinkel, C. A. Styles and G. R. Fink, 1985 Ty elements transpose through an RNA intermediate. Cell 40: 491–500.

Brachat, A., J. V. Kilmartin, A. Wach and P. Philippsen, 1998 Saccharomyces cerevisiae cells with defective spindle pole body outer plaques accomplish nuclear migration via half-bridge-organized microtubules. Mol Biol Cell 9: 977–991.

Bylund, G. O., and P. M. Burgers, 2005 Replication protein A-directed unloading of PCNA by the Ctf18 cohesion establishment complex. Mol Cell Biol 25: 5445–5455.

Carminati, J. L., and T. Stearns, 1997 Microtubules orient the mitotic spindle in yeast through dynein-dependent interactions with the cell cortex. J Cell Biol 138: 629–641.

Chang, M., M. Bellaoui, C. Boone and G. W. Brown, 2002 A genome-wide screen for methyl methanesulfonate-sensitive mutants reveals genes required for S phase progression in the presence of DNA damage. Proc Natl Acad Sci U S A 99: 16934–16939.

Costanzo, M., B. Vandersluis, E. N. Koch, A. Baryshnikova, C. Pons et al., 2016 A global genetic interaction network maps a wiring diagram of cellular function. Science 353.

Cottingham, F. R., and M. A. Hoyt, 1997 Mitotic spindle positioning in Saccharomyces cerevisiae is accomplished by antagonistically acting microtubule motor proteins. J Cell Biol 138: 1041–1053.

Dubarry, M., C. Lawless, A. P. Banks, S. Cockell and D. Lydall, 2015 Genetic Networks Required to Coordinate Chromosome Replication by DNA Polymerases alpha, delta, and epsilon in Saccharomyces cerevisiae. G3 (Bethesda) 5: 2187–2197.

Escusa, S., J. Camblong, J. M. Galan, B. Pinson and B. Daignan-Fornier, 2006 Proteasome- and SCF-dependent degradation of yeast adenine deaminase upon transition from proliferation to quiescence requires a new F-box protein named Saf1p. Mol Microbiol 60: 1014–1025.

Escusa, S., D. Laporte, A. Massoni, H. Boucherie, A. Dautant et al., 2007 Skp1-Cullin-F-box-dependent degradation of Aah1p requires its interaction with the F-box protein Saf1p. J Biol Chem 282: 20097–20103.

Fillingham, J., J. Recht, A. C. Silva, B. Suter, A. Emili et al., 2008 Chaperone control of the activity and specificity of the histone H3 acetyltransferase Rtt109. Mol Cell Biol 28: 4342–4353.

Finley, D., E. Ozkaynak and A. Varshavsky, 1987 The yeast polyubiquitin gene is essential for resistance to high temperatures, starvation, and other stresses. Cell 48: 1035–1046.

Garfinkel, D. J., J. D. Boeke and G. R. Fink, 1985 Ty element transposition: reverse transcriptase and virus-like particles. Cell 42: 507–517.

Gaupel, A. C., T. Begley and M. Tenniswood, 2014 High throughput screening identifies modulators of histone deacetylase inhibitors. BMC Genomics 15: 528.

Gellon, L., D. F. Razidlo, O. Gleeson, L. Verra, D. Schulz et al., 2011 New functions of Ctf18-RFC in preserving genome stability outside its role in sister chromatid cohesion. PLoS Genet 7: e1001298.

Giaever, G., A. M. Chu, L. Ni, C. Connelly, L. Riles et al., 2002 Functional profiling of the Saccharomyces cerevisiae genome. Nature 418: 387–391.

Hartman, J. L. T., and N. P. Tippery, 2004 Systematic quantification of gene interactions by phenotypic array analysis. Genome Biol 5: R49.

Huffaker, T. C., J. H. Thomas and D. Botstein, 1988 Diverse effects of beta-tubulin mutations on microtubule formation and function. J Cell Biol 106: 1997–2010.

Mayer, M. L., I. Pot, M. Chang, H. Xu, V. Aneliunas et al., 2004 Identification of protein complexes required for efficient sister chromatid cohesion. Mol Biol Cell 15: 1736–1745.

Mclellan, J., N. O’Neil, S. Tarailo, J. Stoepel, J. Bryan et al., 2009 Synthetic lethal genetic interactions that decrease somatic cell proliferation in Caenorhabditis elegans identify the alternative RFC CTF18 as a candidate cancer drug target. Mol Biol Cell 20: 5306–5313.

Merkle, C. J., L. M. Karnitz, J. T. Henry-Sanchez and J. Chen, 2003 Cloning and characterization of hCTF18, hCTF8, and hDCC1. Human homologs of a Saccharomyces cerevisiae complex involved in sister chromatid cohesion establishment. J Biol Chem 278: 30051–30056.

Okimoto, H., S. Tanaka, H. Araki, E. Ohashi and T. Tsurimoto, 2016 Conserved interaction of Ctf18-RFC with DNA polymerase epsilon is critical for maintenance of genome stability in Saccharomyces cerevisiae. Genes Cells 21: 482–491.

Palmer, R. E., D. S. Sullivan, T. Huffaker and D. Koshland, 1992 Role of astral microtubules and actin in spindle orientation and migration in the budding yeast, Saccharomyces cerevisiae. J Cell Biol 119: 583–593.

Parsons, A. B., R. L. Brost, H. Ding, Z. Li, C. Zhang et al., 2004 Integration of chemical-genetic and genetic interaction data links bioactive compounds to cellular target pathways. Nat Biotechnol 22: 62–69.

Petronczki, M., B. Chwalla, M. F. Siomos, S. Yokobayashi, W. Helmhart et al., 2004 Sister-chromatid cohesion mediated by the alternative RF-CCtf18/Dcc1/Ctf8, the helicase Chl1 and the polymerase-alpha-associated protein Ctf4 is essential for chromatid disjunction during meiosis II. J Cell Sci 117: 3547–3559.

Scholes, D. T., M. Banerjee, B. Bowen and M. J. Curcio, 2001 Multiple regulators of Ty1 transposition in Saccharomyces cerevisiae have conserved roles in genome maintenance. Genetics 159: 1449–1465.

Shiomi, Y., C. Masutani, F. Hanaoka, H. Kimura and T. Tsurimoto, 2007 A second proliferating cell nuclear antigen loader complex, Ctf18-replication factor C, stimulates DNA polymerase eta activity. J Biol Chem 282: 20906–20914.

Spencer, F., S. L. Gerring, C. Connelly and P. Hieter, 1990 Mitotic chromosome transmission fidelity mutants in Saccharomyces cerevisiae. Genetics 124: 237–249.

Watanabe, M., D. Watanabe, S. Nogami, S. Morishita and Y. Ohya, 2009 Comprehensive and quantitative analysis of yeast deletion mutants defective in apical and isotropic bud growth. Curr Genet 55: 365–380.

Zhao, R., M. Davey, Y. C. Hsu, P. Kaplanek, A. Tong et al., 2005 Navigating the chaperone network: an integrative map of physical and genetic interactions mediated by the hsp90 chaperone. Cell 120: 715–727.

